# Octadecaneuropeptide prevents toxicity induced by 6-hydroxydopamine in cultured rat astrocyte: involvement of the endogenous antioxidant systems and the intrinsic apoptotic pathway

**DOI:** 10.1101/266379

**Authors:** Hadhemi Kaddour, Yosra Hamdi, David Vaudry, Jérôme Leprince, Hubert Vaudry, Marie-Christine Tonon, Mohamed Amri, Olfa Masmoudi-Kouki

## Abstract

Oxidative stress, associated with various neurodegenerative diseases, induces imbalance in ROS generation, impairs cellular antioxidant defences and finally triggers both neurons and astroglial cell death by apoptosis. Astrocytes specifically synthesize and release endozepines, a family of regulatory peptides, including the octadecaneuropeptide (ODN). We have previously reported that ODN is a potent neuroprotective agent that prevents 6-OHDA-induced apoptotic neuronal death. The purpose of the present study was to investigate the potential glioprotective effect of ODN on 6-OHDA-induced oxidative stress and cell death in cultured rat astrocytes. Incubation of astrocytes with graded concentrations of ODN (10^−14^ to 10^−8^ M) inhibited 6-OHDA-evoked cell death in a concentration- and time-dependent manner. In addition, ODN prevented the decrease of mitochondrial activity and caspase-3 activation induced by 6-OHDA. Toxin-treated cells exhibited high level of ROS associated with a generation of H_2_O_2_ and O_2_^°-^and a reduction of both SOD and catalase activities. Co-treatment of astrocytes with low concentrations of ODN dose dependently blocked 6-OHDA-evoked ROS production and inhibition of antioxidant enzymes activities. Taken together, these data demonstrate that ODN is a potent glioprotective agent that prevents 6-OHDA-induced oxidative stress and apoptotic cell death. ODN is thus a potential candidate to delay neuronal damages in various pathological conditions involving oxidative neurodegeneration.

## Introduction

Astrocytes have long been considered as just providing trophic support for neurons, but recently many reports have demonstrated their importance in many functions such as neurotransmission, cell signaling, inflammation, and synapse modulation (Ricci et al., 2009) (Parpura et al., 2012). Currently, involvement of astrocytes dysfunction in the pathophysiology of neurological disorders, including neurodegenerative disease and stroke was highlighted. Parkinson’s disease (PD) is a progressive neurodegenerative disorder that is primarily characterized by a progressive loss of dopaminergic neurons (Henning et al., 2008). One of the widely used models of PD involves treatment with 6-hydroxydopamine (6-OHDA), a neurotoxin that causes selective damage of catecholaminergic neurons both in vivo and in vitro (Blesa and Przedborski, 2014). In vitro experiments shows that the sensitivity of neurons to 6-OHDA has been mainly associated with an auto-oxidation-mediated overproducing of reactive oxygen species (ROS) such as hydrogen peroxide (H_2_O_2_), superoxide anion (O_2_^°-^) and hydroxyl radical (Han et al., 2014; Soto-Otero et al., 2000). The main cellular defense mechanism to cope with oxidative stress is the antioxidant molecules and enzymes (glutathione, catalase, superoxide dismutase and glutathione peroxidase), which play a major neuroprotective role against the deleterious effects of 6-OHDA (Kiasalari et al., 2015; Wu et al., 2015). Treatment of neuroblastoma cell lines with astrocyte conditioned medium significantly increased resistance to oxidative stress and thus neuronal survival against 6-OHDA. In particular, it has been shown that administration of astrocyte conditioned medium provoked a reduction of neuron death and a significant stimulation of glutathione peroxidase expression and glutathione levels (Gardaneh et al., 2011; Gharib et al., 2013). These data highlight the role of astroglial cells to maintain cerebral antioxidant competence and to prevent neuronal damage.

Activation of glia is a common pathological feature of multiple neurodegenerative conditions, which is characterized by a series of metabolic and morphological changes of astrocytes (Lev et al., 2013; Ji et al., 2008). Reactive astrogliosis, abundantly occurred in brain insults (Burda et al., 2014), are found in the substantia nigra and cerebral cortex of Parkinson’s disease brains as well as in 6-OHDA-treated Parkinson’s disease animal models (Chen et al., 2005; Gomide et al., 2005). Although activated astrocytes were traditionally thought to impede neuronal regeneration by forming glial scars (Bortolanza et al., 2014), growing evidence has indicated that reactive astrocytes contribute by crucial benefits in functional recovery of brain injuries (Gomide et al., 2005; L’Episcopo et al., 2011; Pekny and Nilsson, 2005). Concurrently, Analysis of post mortem PD brains has revealed an extensive neuronal loss within areas poorly populated with astrocytes (Damier et al., 1993; Mirza et al., 2000). Moreover, it has been shown that, in rodents, injection of 6-OHDA within the nigrostriatal pathway produced loss of astrocytes associated with the degeneration of dopaminergic neurons (Espinosa-Oliva et al., 2014). Indeed, Astroglial cells are the source of neurotrophic factors that can protect remaining neurons against oxidative stress by detoxifying ROS through the antioxidant enzymes and ROS scavenger molecules (Nakagawa and Schwartz, 2004; Gardaneh et al., 2011; Nakagawa et al., 2005; Takuma et al., 2004). Although glial cells are equipped with high levels of protective and antioxidant compounds, astrocyte death has been demonstrated in human and animal models of brain injuries (Lim et al., 2014). As, loss of glial cells may strongly affect neuronal survival, protection of astrocytes from oxidative insult appears essential to maintain brain function.

The octadecaneuropeptide (ODN) is a peptide generated through the proteolytic cleavage of the 86-amino acid precursor diazepam-binding inhibitor (DBI) (Guidotti et al., 1983) which is exclusively expressed in astroglial cells in the central nervous system (CNS) of mammals (Tonon MC, 2013). DBI and its derived peptides, including ODN, are collectively termed endozepines (Tonon MC, 2013). The primary structure of ODN has been strongly preserved during evolution, suggesting that this peptide exerts a wide range of biological functions such as the control of food intake, sleep, aggressiveness and anxiety (Tonon MC, 2013). At the cellular level, numerous data indicate that ODN acts both as an autocrine and paracrine factor modulating proliferation and differentiation of astroglial and neuron cells, respectively (Alfonso et al., 2012; Gandolfo et al., 1999). In addition, we have recently shown that ODN exerts a potent protective effect upon the deleterious action of oxidative stress in cultured astrocytes (Hamdi et al., 2012b) and cerebellar granule neurons (Kaddour et al., 2013). These data suggest that the endozepine ODN may be implicated, as an endogenous actor, in the neuroprotective and antioxidant proprieties of astroglial cells.

It is well established that 6-OHDA provokes cell apoptosis through the production of ROS leading to apoptosis and activation of caspase-3 in both neurons and glial cells (Raicevic et al., 2005; Hernandez-Baltazar et al., 2013). Since, the gliopeptide ODN protects granule neurons against apoptosis induced by 6-OHDA exposure (Kaddour et al., 2013), we hypothesize that ODN may also protect astroglial cell against 6-OHDA oxidative assault. Therefore, the purpose of the present study was to examine the potential protective action of ODN against 6-OHDA-induced cell death and to investigate the effects of the peptide on some oxidative stress-related parameters in cultured rat astrocyte.

## Materials and Methods

### Animals

Wistar rats (Pasteur Institute, Tunis, Tunisia, and Charles River Laboratories, St Germain sur l’Arbresle, France) were kept in a temperature-controlled room (21 ± 1°C) under an established photoperiod (lights on from 7:00 am to 7:00 pm) with free access to food and water. Experiments were performed in accordance with American Veterinary Medical Association. Approval for these experiments was obtained from the Medical Ethical Committee For the Care and Use of Laboratory Animals of Pasteur Institute of Tunis. Approval Nu FST/LNFP/Pro152012.

### Chemicals

Dulbecco’s modified Eagle’s medium (DMEM), D(+)-glucose, F-12, L-glutamine, foetal bovine serum (FBS), *N*-2-hydroxyethylpiperazine-*N*-2-ethane sulfonic acid buffer solution (HEPES), antibiotic antimycotic solution and trypsin-EDTA were purchased from Gibco (Invitrogen, Grand Island NY, USA). 6-OHDA, fluorescein diacetate-acetoxymethyl ester (FDA-AM), bovine serum albumin (BSA), nitroblue tetrazolium (NBT), Triton X-100, bovine liver catalase, DL-epinephrine, dimethysulfoxyde (DMSO), dihydrorhodamine 123 (DHR123) and insulin were obtained from Sigma Aldrich (St. Louis, MO, USA). The LDH kit assay was commercialized by Bio-Maghreb (Tunis, Tunisia). 5-6-chloromethyl-2′,7′-dichlorodihydrofluorescein diacetate, acetyl ester (CM-H2DCFDA), 5,5”,6,6”-tetrachloro-1,1”,3,3”-tetraethylbenzimidazolylcarbocyanine iodide (JC-1) and dihydroethidium (DHE) were from Molecular Probes (Eugene, Oregon, USA). The Apo-ONE homogeneous caspase-3/7 assay kit was purchased from Promega (Charbonnières, France). Rat ODN (QATVGDVNTDRPGLLDLK) was synthesized by using the standard fluorenylmethyloxycarbonyl (Fmoc) procedure, as previously described (Leprince et al., 2001).

### Secondary culture of cortical rat astrocytes

Secondary cultures of rat cortical astrocytes were prepared from 1- or 2-day-old Wistar rats as previously described (Hamdi et al., 2011) with minor modifications. Briefly, cerebral hemispheres were collected in DMEM/F12 (2:1; v/v) culture medium supplemented with 2 mM glutamine, 1‰ insulin, 5 mM HEPES, 0.4% glucose and 1% of the antibiotic-antimycotic solution. The tissues were dissociated mechanically with a syringe equipped with a 1-mm gauge needle, and filtered through a 100-μm sieve (Falcon, Franklin Lakes, NJ, USA). Dissociated cells were resuspended in culture medium supplemented with 10% FBS, plated in 175-cm^2^ flasks (Greiner Bio-one GmbH, Frickenhausen, Germany) and incubated at 37°C in a 5% CO_2_/95% O_2_ atmosphere. When cultures were confluent, astrocytes were isolated by shaking overnight the flasks with an orbital agitator. Adhesive cells were detached by trypsination and preplated for 5 min to discard contaminating microglial cells. Then, the non-adhering astrocytes were harvested and plated on 35-mm Petri dishes at a density of 0.3 × 10^6^ cells/ml and in 24-well plates at a density of 8 × 10^4^ cells/ml. After 5 days (DIV5), more than 99% of the cells were labeled with antibodies against glial fibrillary acidic protein (Castel et al., 2006). All experiments were performed on 5- to 7-day-old secondary cultures.

### Cell cytotoxicity measurement

Cultured astrocytes were incubated at 37°C with fresh serum-free culture medium in the absence or presence of 6-OHDA and/or ODN for different times. At the end of the incubation, the cytotoxicity of 6-OHDA on astrocytes was determined by measurement of LDH activity in culture medium. The amount of LDH released into medium was measured by LDH assay kit (Bio-Maghreb, Ariana, Tunisia) according to the manufacturer’s instructions. The results were expressed as a percentage of total LDH release after cell lysis with 1% Triton X-100 in saline phosphate buffer (PBS, 0.1 M, pH 7.4).

### Cell survival measurement

The surviving of cells was quantified by measuring FDA in cultured astrocytes. Cells seeded into 24-well plates were subjected to various treatments, incubated in the dark with FDA-AM (15 µg/mL, 8 min), rinsed twice with PBS, and lysed with a 10 mM Tris-HCl solution containing 1% sodium dodecyl sulfate (SDS). Fluorescence intensity (λ excitation = 485 nm and λ emission = 538 nm) was measured with a FL800TBI fluorescence microplate reader (Bio-Tek Instruments, Winooski, VT, USA).

### Mitochondrial activity

Mitochondrial membrane potential was quantified using the JC-1 probe. Cells seeded into 24-well plates were incubated at 37°C for 72 h with fresh serum-free culture medium in the absence or presence of 6-OHDA and/or ODN. At the end of the experiments, astrocytes were treated for 15 min with the JC-1 probe (10 µg/mL) and then washed twice with PBS. Fluorescence intensity was measured with a FL800TBI fluorescence microplate reader and expressed as a ratio of the fluorescence emission at 590 nm (orange, intact mitochondrial membrane potential) versus 530 nm (green, collapsed mitochondrial membrane potential).

### Caspase 3 activity measurement

The effect of ODN on 6-OHDA-induced increase of caspase 3 activity was measured by using the Apo-ONE Homogeneous Caspase-3/7 kit (Promega). At the end of the experiments, cells were washed twice with PBS and resuspended DMEM (100 μL) mixed with 1X caspase assay buffer containing 25 μM caspase-3 substrate. Caspase-3 activity was calculated from the slope of the fluorescence measured every 15 min for 3 h with excitation at 485 nm and emission at 530 nm, and expressed as a percentage of the control.

### Intracellular reactive oxygen species measurement

Reactive oxygen species were detected by measuring the fluorescence of 2′,7′-dichlorodihydrofluorescein (DCF) resulting from the deacetylation and oxidation of the non-fluorescent compound DCFH2-DA. Cells seeded into 24-well plates were exposed to 6-OHDA with or without ODN, incubated with 10 μM cell-permeant DCFH_2_-DA in serum-free loading medium for 45 min at 37°C and then washed twice with PBS. DCF fluorescence (λ excitation = 485 nm and λ emission = 538 nm) was measured with a FL800TBI fluorescence microplate reader.

### Hydrogen peroxide measurement

H_2_O_2_ generation were detected by using the dihydrorhodamine 123 (DHR123) which is an uncharged and nonfluorescent fluorochrome that can passively diffuse across membranes where it is oxidized to cationic rhodamine 123 which localizes in the mitochondria and exhibits green fluorescence. After treatments with substances of interest, astrocytes were loaded with 6 µM DHR123 for 15 min. Excess of DHR123 was removed by washing with PBS. Fluorescence intensity measurement (λ excitation = 488 nm and λ emission = 520 nm) was measured using a microplate reader (Bio-Tek FL800TBI).

### Intracellular superoxide anion measurement

O_2_^°-^ levels were detected by measuring the fluorescence of ethidium which derived from the oxidation of the non-fluorescent compound dihydroethidium (DHE). Glial cells were incubated at 37 °C with fresh serum-free medium in the absence or presence of test substances. At the end of the incubation, the cells were incubated for 15 min in fresh medium containing 2 µM DHE and then washed twice with PBS. Ethidium fluorescence (λ excitation = 488 nm and λ emission = 575 nm) was measured with a microplate reader (Bio-Tek FL800TBI).

### Respiratory burst assay

The formation of the O_2_°^−^, produced by respiratory burst, was assessed by the reduction of NBT into a blue precipitate inside the cell. At the end of the incubation with test substances, cultured astrocytes were washed once with PBS at 37°C, and then treated with NBT (1mg/ml), in PBS containing BSA (1mg/ml) for 2h. At the end of the incubation, cells were examined on an inverted microscope (Nikon) to visualize the formation of blue crystals.

### Measurement of antioxidant enzyme activities

Cultured cells were incubated at 37°C for 72 h with fresh serum-free medium in the absence or presence of test substances. At the end of the incubation, cells were washed twice with PBS and total cellular proteins were extracted by using the lysis buffer containing 50 mM Tris– HCl (pH 8), 10 mM EDTA, 100 µM phenylmethyl-sulfonylfluoride and 1% Triton X-100. The homogenate was centrifuged (16, 000 g, 4°C, 20 min) and the cellular extract contained in the supernatant was stored at ^_^20°C until enzyme activity determinations.

SOD activity was measured using a spectrophotometric assay, which consists in measuring epinephrine autoxidation induced by superoxide anion. Samples, prepared as described above, were incubated for 3 min with a mixture containing bovine catalase (0.4 U/μL), DL-epinephrine (5 mg/mL) and Na_2_CO_3_ / NaHCO_3_ buffer (62.5 mM, pH 10.2). The oxidation of epinephrine was measured at 480 nm with a Bio-Rad spectrophotometer (Bio-Rad Laboratories, Philadelphia, USA).

Catalase activity was determined on the basis of the decrease of H_2_O_2_. Samples, prepared as described above, were mixed with 30 mM H_2_O_2_ in PBS. The disappearance of H_2_O_2_ was measured at 240 nm for 180 s at 30 s intervals. Catalase activity was calculated using the extinction coefficient of 40 mM-1 cm−1 for H_2_O_2_.

### Statistical analysis

Data are expressed as the mean ± SEM from three independent experiments. Statistical analysis of the data was performed by using ANOVA, followed by Bonferroni’s test. A *p* value of 0.05 or less was considered as statistically significant.

## Results

### Glioprotective effect of ODN against 6-OHDA-induced astroglial cell death

Time courses experiments revealed that incubation of astrocytes with 6-OHDA (120 µM) induced a time-dependent decrease of the proportion of surviving cells associated with an increase of LDH levels in the culture medium (Fig. 1A). Furthermore, treatment of astroglial cells during 72 h with graded concentrations of 6-OHDA (10 to 200 µM) provoked a dose-dependent decrease of living cells and increase of cell death (Fig. 1B). The concentration of 120 µM 6- OHDA, which killed 50% of the cells after 72 h of treatment, was used in all subsequent experiments. In contrast, co-incubation of cells with 6-OHDA (120 µM) and graded concentrations of ODN (10^−14^ to 10^−8^M) for 72 h provoked a biphasic mirror effect on astroglial survival and LDH release (Fig. 1C). The protective action of ODN was concentration-dependent ranging from (10^−11^ to 10^−8^M) and complete reversal of the toxic effect of 6-OHDA was obtained with 10^−10^ M and 10^−9^ M ODN (Fig. 1C). It’s important to note that incubation with ODN alone did not affect astrocytes survival whatever the time or the dose.

**Fig. 1.**
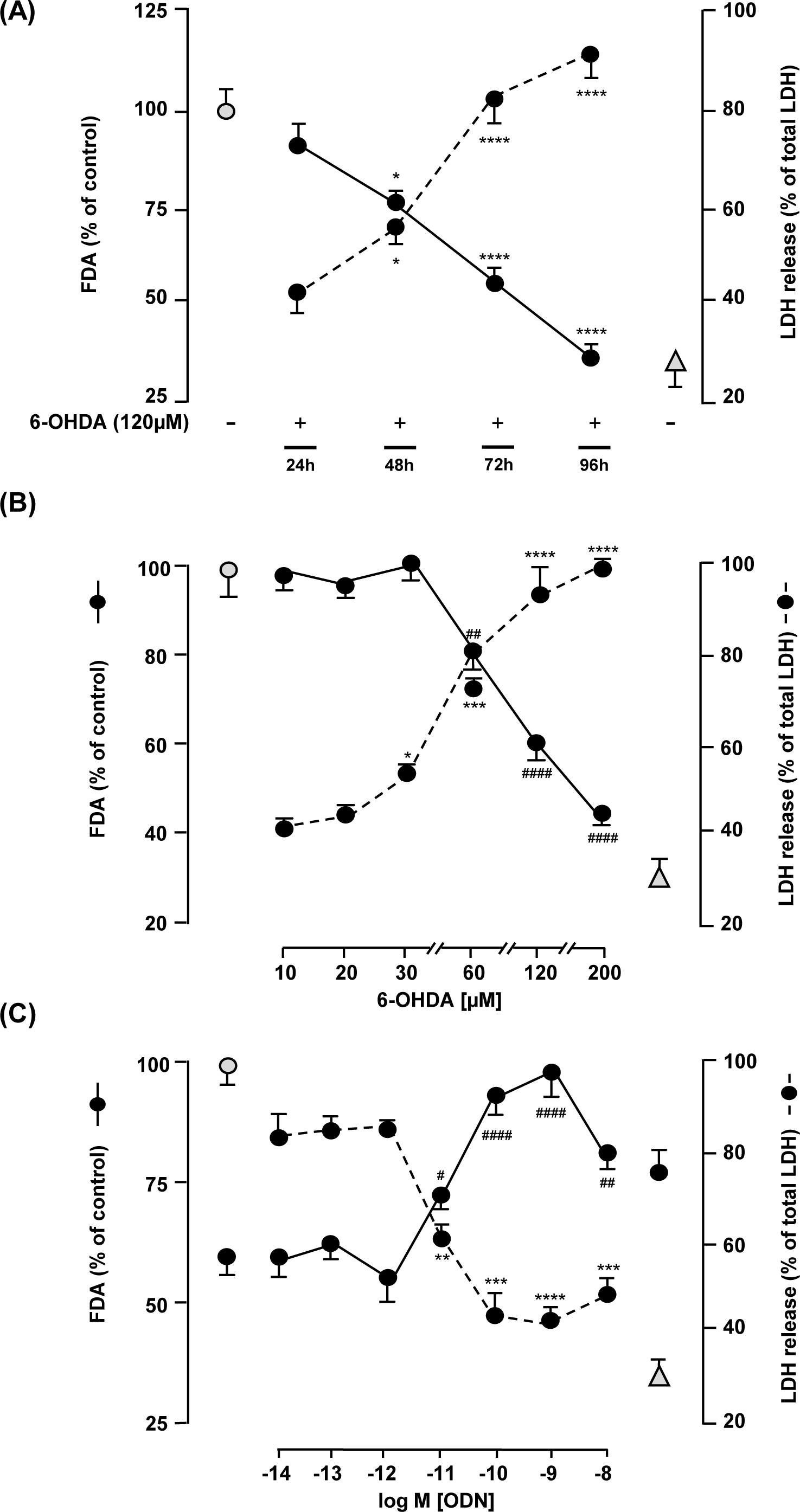
Glioprotective effect of ODN on 6-OHDA-induced cultured rat astrocytes death. (A) Effect of 6-OHDA (120 µM) on astrocytes survival after 24 h, 48 h, 72 h and 96 h of treatment. (B) Effect of graded concentrations of 6-OHDA (10 – 200 µM) on astrocytes survival and LDH release after 72 h. (C) Effect of graded concentrations of ODN (10^−14^ to 10^−8^ M) on 6-OHDA (120 μM) induced astrocytes death after 72 h of treatment. Cell survival (^___^) was quantified by measuring FDA fluorescence intensity, and the results are expressed as percentages of the control (grey circle). Cell death (- - -) was determined by measuring of LDH activity in culture medium, and the results are expressed as percentages of total LDH released in Triton-lysed cells (grey triangle). Each value is the mean (± SEM) from at least 6 different wells from 3 independent cultures. ANOVA followed by Bonferroni’s test: ^*^*p* < 0.05; ^**^*p* < 0.01; ^***^*p* < 0.001; ^****^*p* < 0.0001 *vs.* control. ^#^*p* < 0.05; ^##^*p* < 0.01; ^####^*p* < 0.0001 *vs.* 6-OHDA-treated cells.

### ODN exerts its glioprotective effect through the intrinsic mitochondrial and caspase-3 pathway

Previous reports have shown that 6-OHDA inhibit the mitochondrial electron transport chain and to induce a syndrome closely resembling PD (Glinka and Youdim, 1995) and cause caspase-3 activation in different type of cells, which leads to typical apoptotic cell death (Kaddour et al., 2013) (Wu et al., 2015). Considering this, we first examined the effect of ODN on the integrity of mitochondria by measuring the membrane potential using the fluorescent ratiometric probe JC-1 (Fig. 2A, 2B). Treatment of astrocytes with graded concentrations of 6-OHDA (10 to 200 µM) for 72 h, induced a dose-dependent decrease of the 590/530 nm ratio, indicating that the mitochondrial integrity was severely altered by the toxin (Fig. 2A). Addition of glioprotective doses of ODN (10^−10^ and 10^−9^M) in the incubation medium totally suppressed the deleterious effect of 6-OHDA on mitochondrial membrane potential (Fig. 2B). To further explore the mechanism involved in the protective action of ODN, we monitored the activity of caspase-3. Incubation of cultured astrocytes with graded concentrations of 6-OHDA (10 to 200 µM) for 72 h provoked a dose-dependent increase of caspase-3 activity (Fig. 2C). Co-treatment of the cells with protective doses of ODN (10^−10^ to 10^−9^M) totally blocked toxin-induced caspase-3 activation (Fig. 2D). For both experiments, non protective doses of ODN such as 10^−12^ M or 10^−8^ M are enable to block or inhibit partially the toxin action on mitochondria alteration and increase of caspas-3 activity (Fig 2B, 2D).

**Fig. 2.**
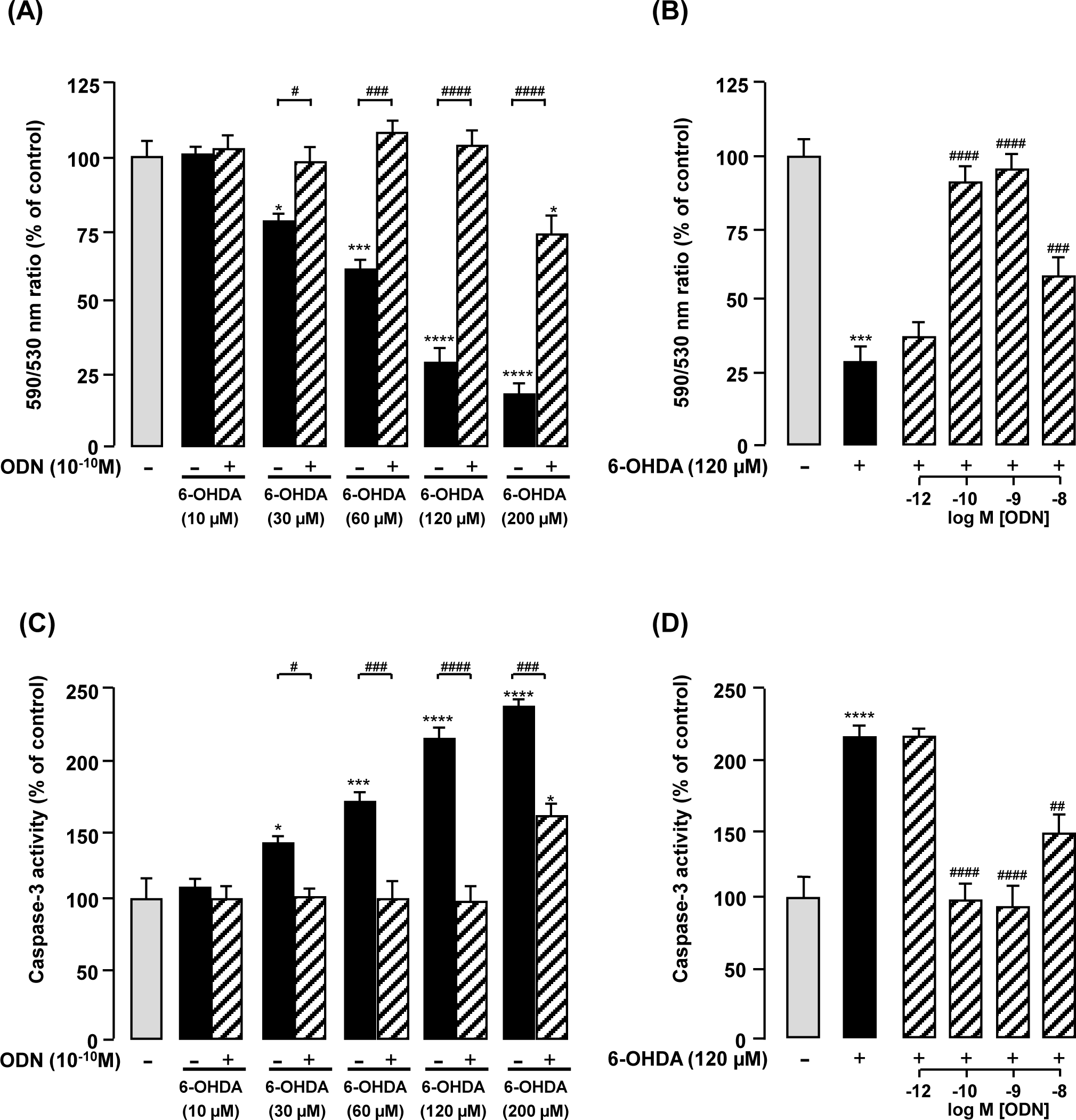
Effect of ODN on 6-OHDA-induced impairment of mitochondrial transmembrane potential and activation of caspase-3 in astrocytes after 72 h of treatment. (A, C) Effect of graded concentrations of 6-OHDA (10 – 200 µM) on mitochondrial transmembrane potential and caspase-3 activity in cells. (B, D) Effect of graded concentrations of ODN (10^−12^ to 10^−8^ M) on 6-OHDA (120 μM) induced impairment of mitochondrial transmembrane potential and activation of caspase-3 in astrocytes. Cells Mitochondrial transmembrane potential was assessed by using the JC-1 probe, and the ratio of fluorescence emissions 590/530 nm was measured as an index of mitochondrial activity. Caspase-3 activity was quantified by measuring the fluorescence of the caspase substrate, Z-DEVD-Rhodamine 110. The results are expressed as percentages of the control. Each value is the mean (± SEM) of at least 3 different wells from 3 independent experiments. ANOVA followed by Bonferroni’s test: ^*^*p* < 0.05; ^***^*p* < 0.001; ^****^*p* < 0.0001 *vs.* control. ^#^*p* < 0.05; ^##^ *p* < 0.01; ^###^*p* < 0.001; ^####^*p* < 0.0001 *vs.* 6-OHDA-treated cells.

### ODN prevents the effect of 6-OHDA on ROS and H_2_O_2_ accumulation

The toxic effect of 6-OHDA has been linked to over-generation of ROS such as H_2_O_2_, O_2_^°-^ and hydroxyl radical (Soto-Otero et al., 2000). To examine whether ODN could block 6-OHDA-induced intracellular ROS accumulation, astrocytes were first labeled with a probe which generates the fluorescent compound dichlorofluorescein (DCF) upon oxidation with ROS. Incubation of cells with graded concentrations of 6-OHDA (10 to 200 µM) for 72 h induced a dose-dependent increase in DCF fluorescence intensity (Fig. 3A). For the dose of interest of 6-ODHA (120 µM), co-incubation of cells with glioprotective doses of ODN (10^−10^ and 10^−9^M) did not modify DCF fluorescence intensity, but totally abolished the effect of 6-OHDA on DCF formation (Fig. 3B). On another hand, we examined the effect of ODN on the formation of H_2_O_2_ by astrocytes. When glial cells were treated with toxic concentrations of 6-OHDA (60 to 200 µM), there were significant increases in DHR123 fluorescence intensity with a maximum increasing at 200 µM (< 120 %; Fig. 3C). Co-treatment of cells with glioprotective doses of ODN (10^−10^ and 10^−9^M) ODN totally suppressed the effect of the toxin on H_2_O_2_production (Fig. 3D).

**Fig. 3.**
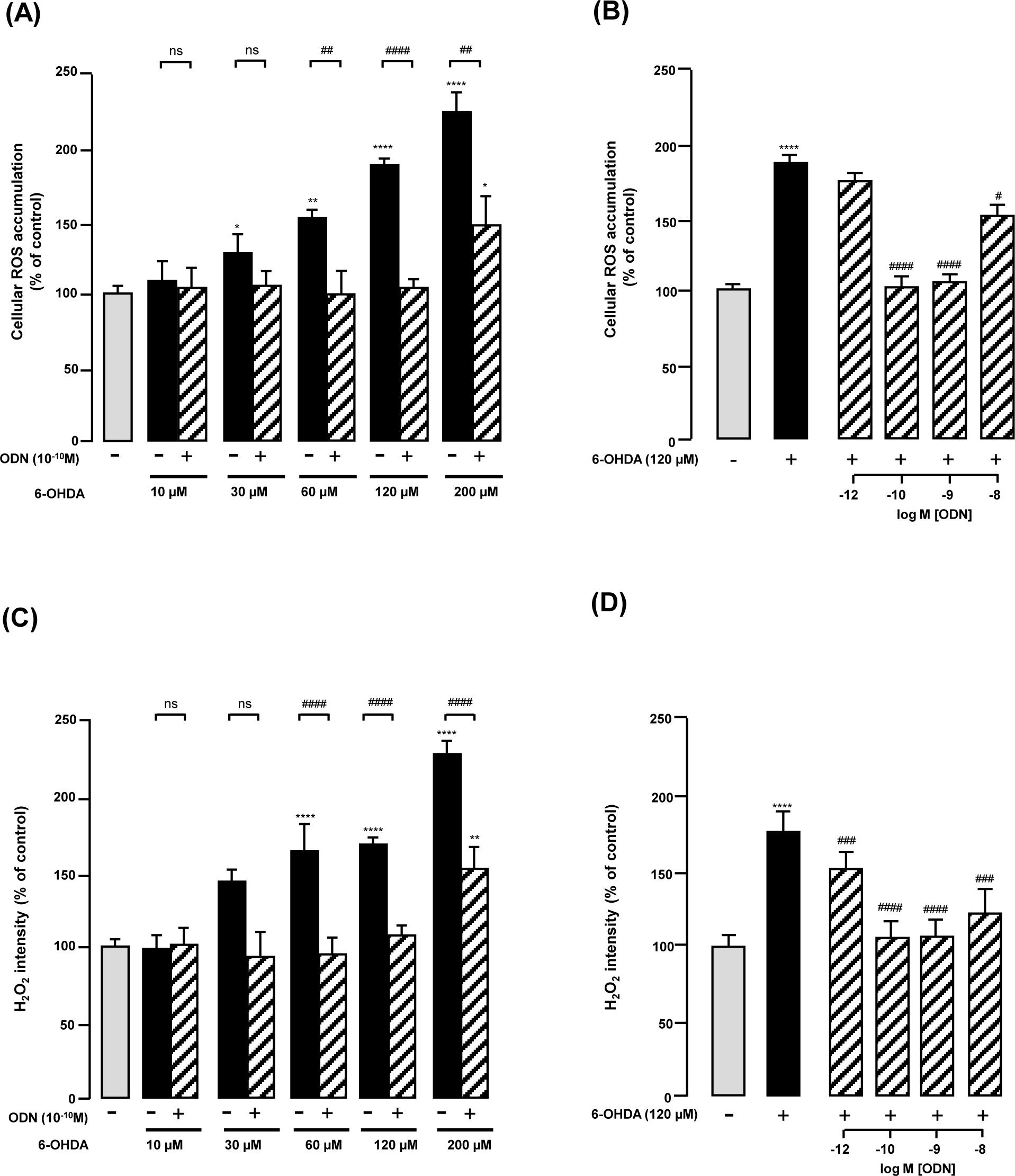
Effect of ODN on 6-OHDA-induced intracellular generation of ROS and accumulation of H_2_O_2_. Cells were incubated for 72 with graded concentration of 6-OHDA (10 – 200 µM) in the absence or presence of ODN (10^−10^M) (A, C). Cells were co incubated with graded concentrations of ODN (10^−12^ to 10^−8^ M) and 6-OHDA (120 μM) (B, D). Cellular ROS and H_2_O_2_generation were quantified by measurement of DCF and DHR 123 fluorescence, respectively The results are expressed as percentages of control. Each value is the mean (± SEM) of at least 3 different wells from 2 independent experiments. ANOVA followed by Bonferroni’s test: ^*^*p* < 0.05;^**^*p* < 0.01; ^****^*p* < 0.0001 vs. control. ^#^*p* < 0.05;^##^ *p* < 0.01; ^###^*p* < 0.001; ^####^*p* < 0.0001; ns, not statistically different vs. 6-OHDA-treated cells.

### ODN prevents the effect of 6-OHDA on superoxide radical generation

Since large amounts of O_2_°^−^ are produced by respiratory burst under a drastic oxidative stress status, we have investigated the effect of ODN on 6-OHDA-induced O_2_°^−^ formation by astroglial cells. Qualitative analysis using the respiratory burst assay shows that control treated astrocytes exhibited very few blue precipitates (reflecting reduction of NBT by O_2_^°-^ production) in the cell bodies (Fig. 4Aa). Treatment of astrocytes with 6-OHDA resulted in a marked increase of the respiratory burst as shown by the labeling of most cell bodies in blue, indicating that mitochondrial integrity was severely impaired by 6-OHDA (Fig. 4Ab). Incubation of cells with ODN suppressed the deleterious effect of 6-OHDA, only few cells exhibited a blue labeling (Fig.4Ac). Quantitative analysis indicated that Incubation of astrocytes with graded concentrations of 6-OHDA (10 to 200 µM) for 72 h provoked a dose-dependent increase in DHE fluorescence intensity (Fig. 4B) and that ODN restored the O_2_^°-^ levels to the control values (Fig. 4B). For the dose of interest of 6-ODHA (120 µM), co-incubation of cells with glioprotective doses of ODN totally abolished the effect of 6-OHDA on DHE formation (Fig.4C).

**Fig. 4.**
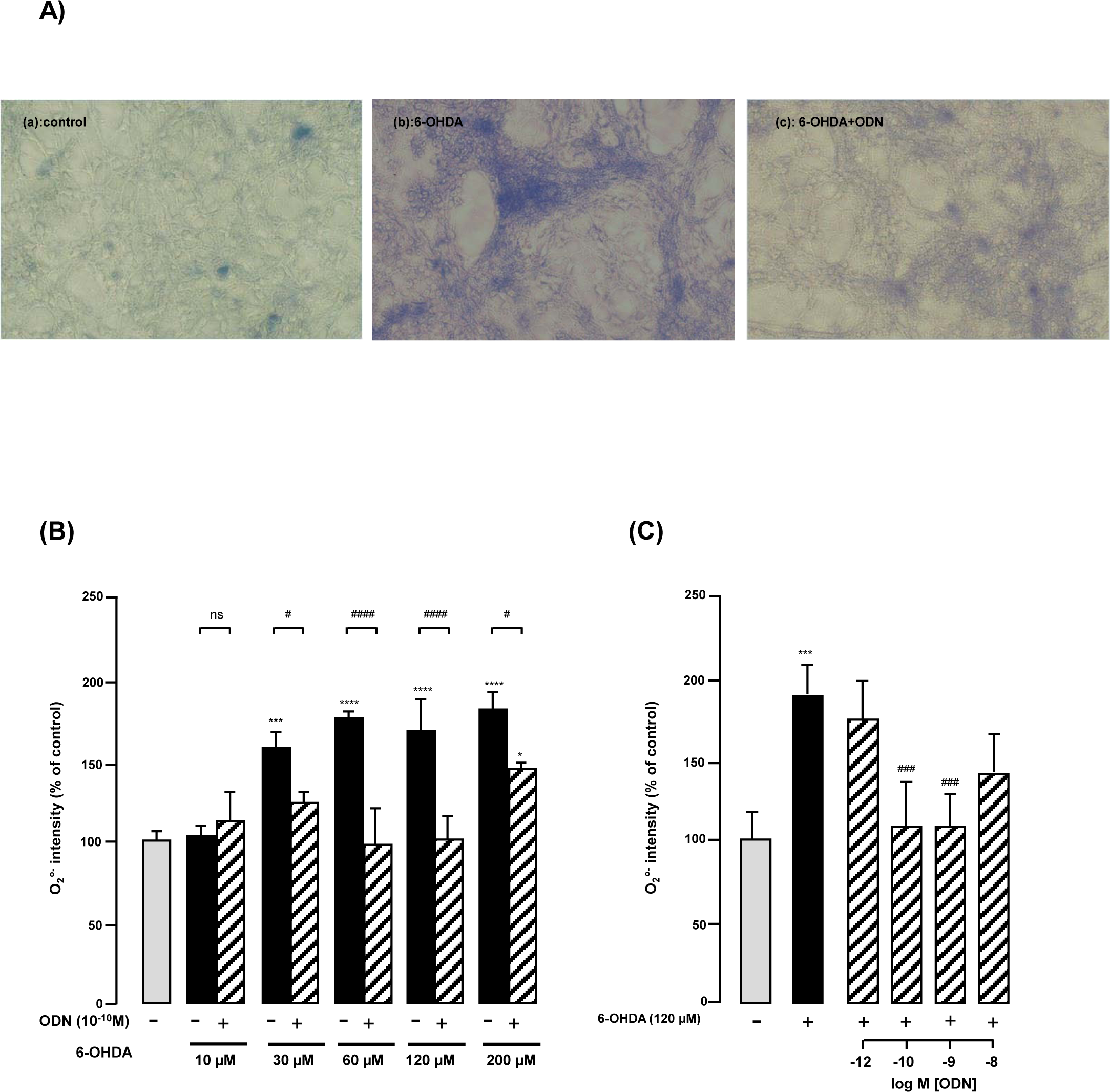
Effect of ODN on 6-OHDA-induced superoxide radical production. (A) Phase-contrast images illustrating the generation of superoxide radicals visualized with the presence of a blue-dark precipitate inside cells. Cells were incubated for 72 h with medium alone (Aa) or with 120 µM 6-OHDA (Ab) alone or in presence of ODN (Ac; 10^−10^ M). Scale bar, 20 μm. (B-C) Cellular superoxide radical levels were quantified by measurement of the fluorescence of oxidized dihydroethidium (DHE). Cells were incubated with increasing concentrations of 6-OHDA (10 – 200 µM) in the presence or absence of ODN (10^−10^ M) (B). Cells were incubated for 72 h with 120 µM 6-OHDA in the absence or presence of graded concentrations of ODN (10^−12^ to 10^−8^ M) (C). The results are expressed as percentage of control. Each value is the mean (± SEM) of at least four different wells from three independent experiments. ANOVA followed by the Bonferroni’s test. ^*^*p* < 0.05;^***^*p* < 0.001; ^****^*p* < 0.0001 vs. control. #*p* < 0.05;^##^ *p* < 0.01; ^####^*p* < 0.0001; ns, not statistically different vs. 6-OHDA-treated cells.

### ODN blocks 6-OHDA-induced inhibition of antioxidant enzyme activities in cultured astrocytes

To further explore the mechanism involved in the protective action of ODN against the oxidative stress generated by 6-OHDA, we monitored the activities of the two antioxidant enzymes SOD and catalase in astrocytes. Exposure of astrocytes with graded concentrations of 6-OHDA (10 to 200 µM) for 72 h, dose-dependently decreased both SOD and catalase activities (Fig. 5A and B). At lower doses, 6-OHDA has no effect on catalase activities (> 60 µM; Fig. 5B). At all concentrations of 6-OHDA tested, ODN restored SOD and catalase activities above control levels. We next examined the effect of graded concentrations of ODN (10^−12^ to 10^−8^M)/6-OHDA (120 µM) co-treatment on enzymes activities. Treatment of glioprotective doses of ODN (10^−10^ to 10^−8^M) in the incubation medium, increased in a dose dependent manner SOD and catalase activities, restoring the levels to the control values (Fig. 5C, 5D).

**Fig. 5.**
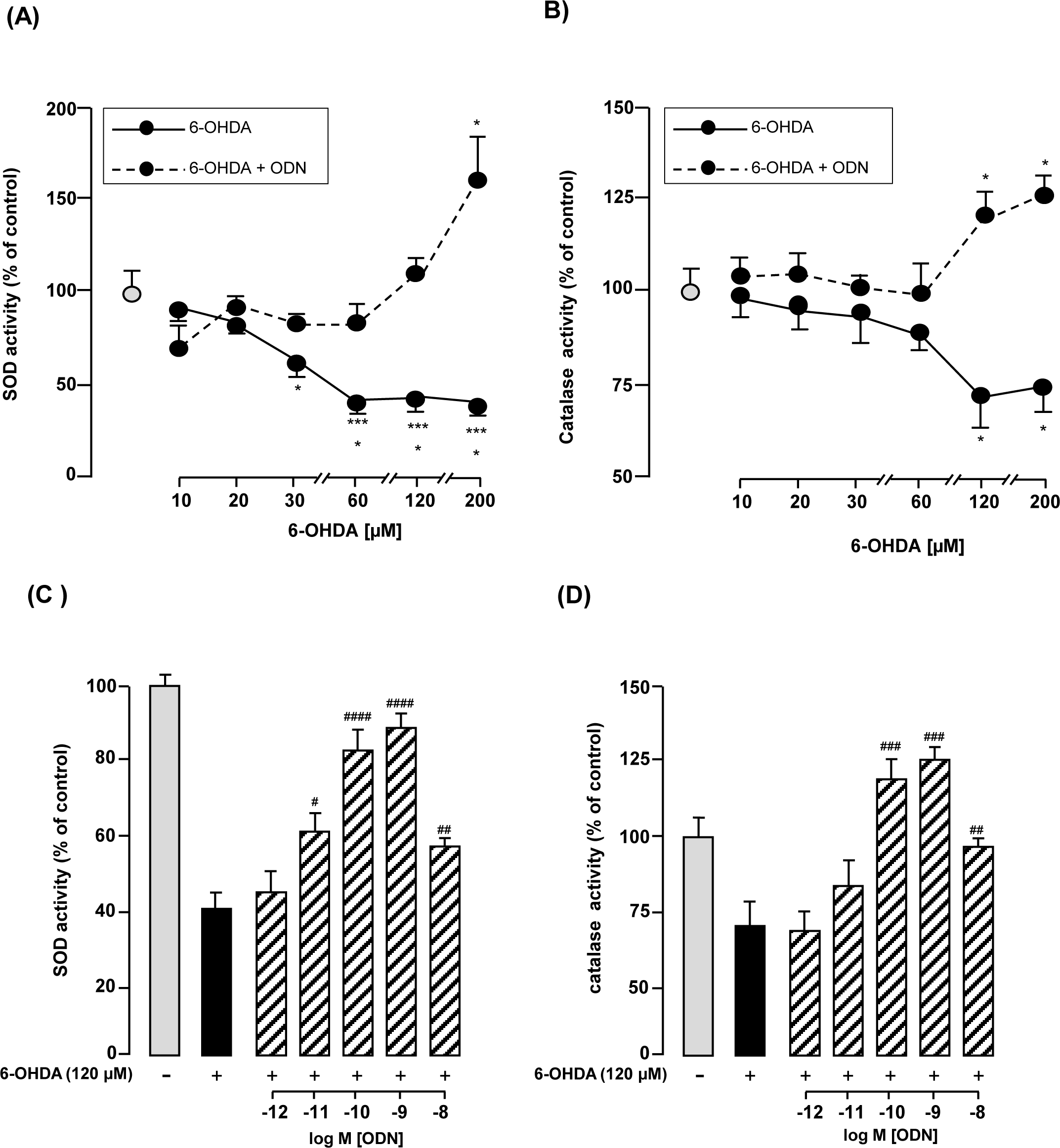
Effect of ODN on modulation of SOD and catalase activities in cultured rat astrocytes. Cells were incubated with increasing concentrations of 6-OHDA (10 – 200 µM) in the presence or absence of ODN (10^−10^ M) (A-B). Cells were incubated for 72 h with 120 µM 6-OHDA in the absence or presence of graded concentrations of ODN (10^−12^ to 10^−8^ M) (C, D). SOD activity was determined by measurement of epinephrine autoxidation induced by superoxide anion, and catalase activity by the decrease of H_2_O_2_. The results are expressed as a percentage of control. Each value is the mean (± SEM) calculated from at least 3 different dishes from 3 independent cultures. ANOVA followed by the Bonferroni’s test. ^*^*p* < 0.05; \*\*\*\**p* < 0.0001 *vs.* control.^#^*p* < 0.05; ^##^ *p* < 0.01; ^###^*p* < 0.001; ^####^*p* < 0.0001 different *vs.* 6-OHDA-treated cells.

## Discussion

It has been previously reported that oxidative stress causes apoptosis in various cell types, including glial cells (Hamdi et al., 2012a). We have previously reported that ODN protects cerebellar granule neurons against 6-OHDA-induced cell death (Kaddour et al., 2013). Here, we demonstrate for the first time that ODN protects cultured rat astrocytes from apoptosis induced by 6-OHDA, and we show that ODN exerts its glioprotective effect through inhibition of ROS generation and caspase-3 activation.

6-OHDA, a neurotoxin classically used to mimicked PD in vivo and in vitro, leads to apoptosis in various cell types that do not express dopaminergic transporters, including C6 astrocytoma cells (Blum et al., 2000; Blum et al., 2001) and rat primary astrocytes (Iwata-Ichikawa et al., 1999; Raicevic et al., 2005). In agreement with this, we observed that exposure of cultured astrocytes to concentrations of 6-OHDA below 60 µM has little effect on astrocyte viability. In contrast at higher concentrations (> 120 µM) and in time-dependent manner (> 48 h), 6-OHDA provoked cell death. This insensitivity of astrocytes at lower doses of the toxin can be explained by their higher capacity for antioxidant defense (Ferrero-Gutierrez et al., 2008) and reports provided evidences that astrocytes were far less sensitive to 6-OHDA toxicity in comparison with neurons (Blum et al., 2000; Kaddour et al., 2013). Moreover, the absence of cell death before 24h can be explain by the fact that injury-induced by the toxin is due to its progressive auto-oxidation in the medium (Hanrott et al., 2006). Incubation of cultured astrocytes with subnanomolar concentrations of ODN (10^−11^ to 10^−8^ M) dose-dependently protects cells against 6-OHDA injury, indicating that ODN is a potent glioprotective agent. While the beneficial effect of ODN against astrocyte cell death is well documented, this is the first report showing that ODN exerts a neuroprotective action against 6-OHDA toxicity.

It is well known that the pathogenesis of age-related diseases such as PD involves the generation of ROS (van Muiswinkel et al., 2004; Hwang, 2013). The mechanism of 6-OHDA toxicity is linked to production of ROS by extracellular auto-oxidation (Hanrott et al., 2006). Thus, in agreement with previous reports, we found that 6-OHDA increases in a concentration-dependent manner the production of ROS in cultured rat astrocytes. The present study reveals that addition of ODN in the culture medium significantly attenuates ROS formation which is likely responsible for a reduction of cell death induced by 6-OHDA. These ROS, induced by 6-OHDA, can easily cross the plasma membrane (Kaczara et al. 2010), react with biological target molecules and damage the mitochondrial membrane (Tian et al., 2008). This damage eventually results in the formation of mitochondrial permeability transition pore leading to release of apoptotic proteins into the cytoplasm and finally to activation of caspase-3, the effectors of apoptotic cell death (Wu et al., 2007). Consistent with other study (Kaddour et al., 2013), cells incubated with 6-OHDA showed a significant reduction of proportion of active mitochondria and a significant activation of caspase-3 activity. Treatment of cells with ODN prevented the deleterious effect of 6-OHDA on mitochondria. Concurrently, ODN markedly suppressed caspase-3 activation induced by the toxin indicating that the protective effect of ODN on astrocytes is attributable to an inhibition of caspase-3 activity through a ROS-mitochondrial-dependent pathway.

Among the ROS that are responsible for oxidative stress, H_2_O_2_ is regarded as a key substance involved in astroglial apoptosis (Ferrero-Gutierrez et al., 2008). In this study, we observed that exposure of cultured astrocytes to 6-OHDA’s concentration of interest (120 µM) induced a significative increase of H_2_O_2_, and O_2_^°-^ intensity above 160 % versus control. O_2_^°-^ can be also converted into H_2_O_2_ by superoxide dismutase leading to the increase of intracellular H_2_O_2_ levels and a self promoting cycle which amplifies H_2_O_2_signaling (Lin and Wang, 2012). Consistent with this observation, it has been shown that astroglial cells express catecholamine transporters, which promote cytosol accumulation of 6-OHDA and H_2_O_2_ generation after deamination by monoamine oxidase of the toxin, indicating that astrocytes might be main target cells for 6-OHDA in the CNS. However, several lines of evidences suggest that ROS-mediated 6-OHDA toxicity is due to formation of highly reactive quinone compounds, H_2_O_2_, O_2_^°-^ and hydroxyl radical via a non enzymatic extracellular auto-oxidation process (Soto-Otero et al., 2000). The present data revealed that ODN blocked the increase of respiration rate and mitochondrial O_2_°- generation induced by 6-OHDA. These data are consistent with previous studies showing that, in cultured astrocytes, ODN induced a rapid stimulation of transcription and activity of antioxidant enzymes which is responsible for detoxification of O_2_^°-^ and H_2_O_2_ (Hamdi et al., 2012b; Hamdi et al., 2015).

Many lines of evidence have proven that disruption of the balance between ROS production and scavenging often lead to apoptotic cell death, which is associated with PD (Shalavadi et al., 2012). It is well established that astroglial cells contain high levels of ROS scavenger molecules such as glutathione and vitamins E and C (Dringen et al., 1999) and antioxidant enzymes Mn- and Cu, Zn-superoxide dismutases (Mn- and Cu, Zn-SOD), catalase and glutathione peroxidase (Lindenau et al., 2000; Sokolova et al., 2001; Saha and Pahan, 2007). Therefore, cellular defenses systems, including antioxidant enzymes have been proposed to be important in protecting cells against ROS (Hamdi et al., 2011). In order to determine the mechanism underlying the protection against ROS conferred by ODN, we examined the antioxidant systems. In agreement with previous reports, we found that 6-OHDA decrease in a concentration dependent manner the activity of SOD and catalase. Co-treatment of astrocytes with subnanomolar concentrations of ODN markedly reduced the inhibitory effect of 6-OHDA on SOD and catalase activities which is likely responsible for a reduction of ROS formation and cell death induced by 6-OHDA. Consistent with this observation, reduction of SOD activity is associated with an exacerbation of oxidative damages in transgenic mouse model of neurodegenerative disease (Schuessel et al., 2005). In addition, exposure of astrocytes to SOD and catalase inhibitors, abrogated the effect of ODN on H_2_O_2_-evoked inhibition of SOD and catalase activities and suppressed the protective effect of ODN against H_2_O_2_-induced astrocyte apoptosis (Hamdi et al., 2011). Altogether, these data indicate that the protective effect of ODN against 6-OHDA- induced cell death in astrocytes is attributable to activation of the antioxidant enzymes that act as scavengers of H_2_O_2_ and ROS.

Several lines of evidence suggest that the protective action of ODN may have a physiopathological significance in neurodegenerative disorders. It is now established that reactive astrocytes contribute to the defense of neurons under moderate oxidative stress by releasing neuroprotective factors. Nevertheless, despite their high antioxidative activities, astrocytes cannot survive and protect neurons under insurmountable oxidative stress (Takuma et al.2004; Giffard and Swanson 2005). Thus, overproduction of the endozepine ODN by astrocyte cells, which acts as a protective agent on astrocytes from oxidative assault, might delay neuronal damages in various pathological conditions involving oxidative neurodegeneration suggesting that ODN might have a therapeutic potential for treatment of cerebral injuries involving oxidative neurodegeneration. A role for ODN on cellular proliferation has been proposed for astrocytes in the central nervous system. In particular, ODN were shown to stimulate cellular division of cultured rat astrocytes (Gandolfo et al., 1999, 2000). Interestingly, Alfonso and colleagues reported that ODN stimulates neurogenesis in adult mouse brain, and that DBI inhibition leads to growth arrest and death of neural progenitors (Alfonso et al., 2012). Altogether, these data suggest that ODN might have a therapeutic potential for treatment of cerebral injuries, by acting as a positive regulator for neuronal survival and differentiation. However, the possible effect of ODN on proliferation andd/or differentiation of neurons deserves better investigations.

